## Supplementary figures and images for "Human hippocampal responses to network intracranial stimulation vary with theta phase"

### Figure 1 - Animation 1

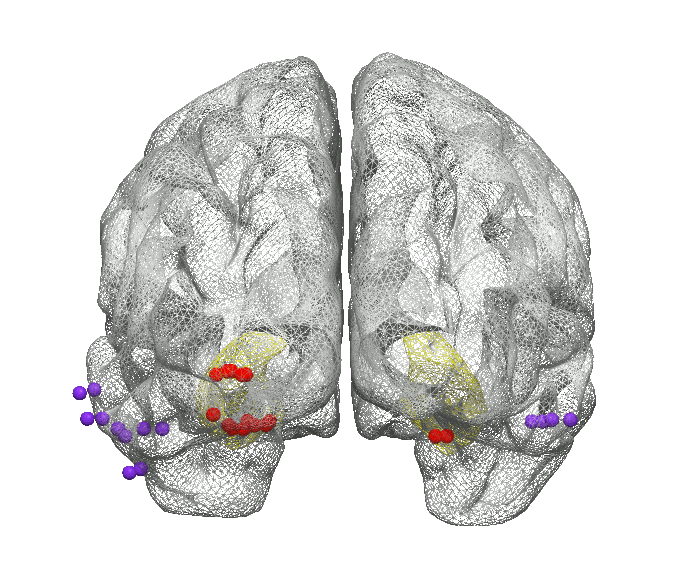
